## Supplementary Figures for "Protein Set Transformer: A protein-based genome language model to power high diversity viromics"

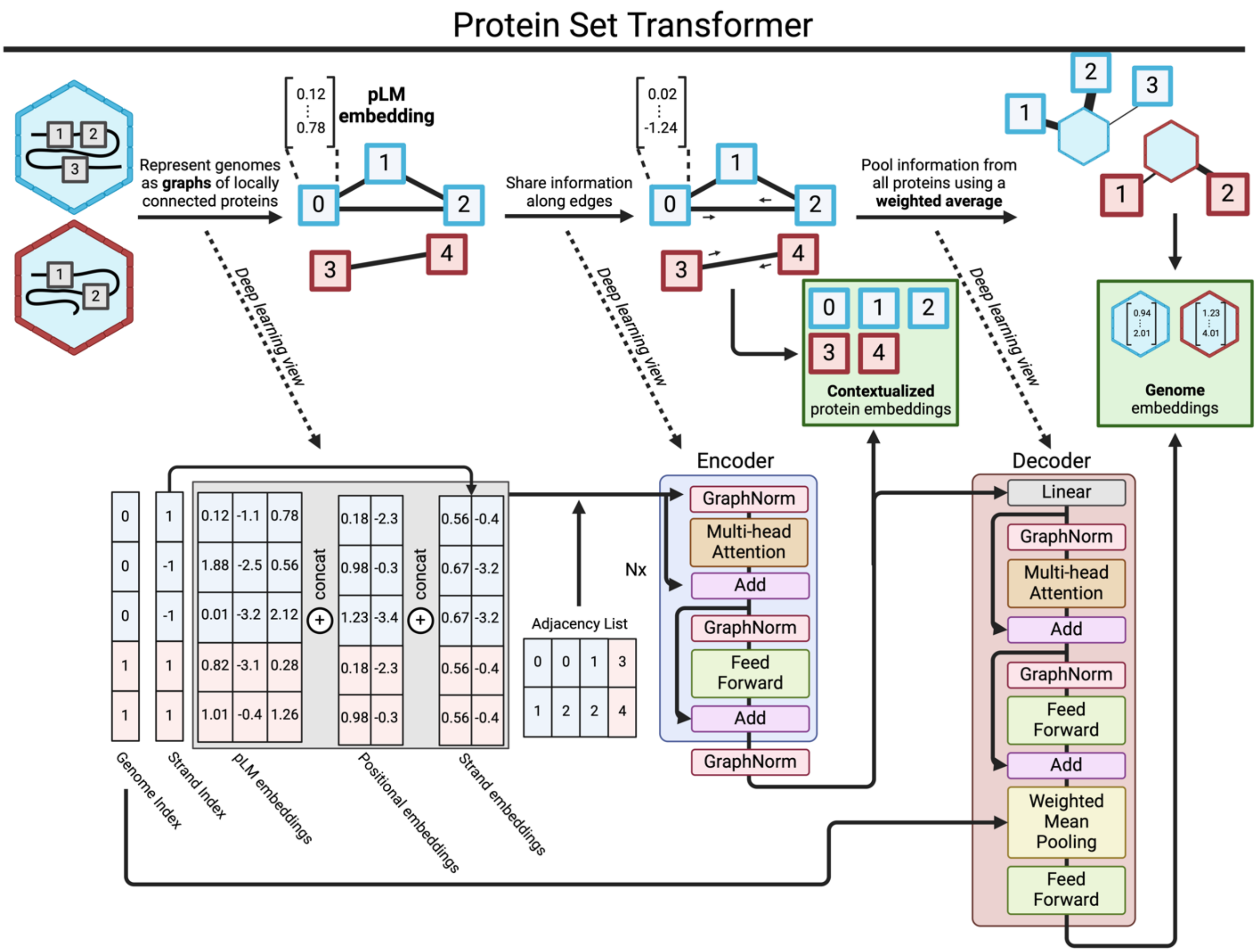


**Supplementary Figure 1**. A machine learning-centric view of the encoder-decoder Protein Set Transformer (PST) architecture. Each genome is internally represented as a graph, composed of subgraph genome chunks whose size is tunable. A minibatch of genomes is represented in a memory-efficient stacked matrix, where the boundaries for the set of proteins from each genome are tracked using indices and offset pointers for efficient random access. At the beginning of training, the ESM2 protein embeddings are concatenated with learnable positional embeddings based on the relative position in each genome and encoding strand embeddings. This is then input to the PST encoder, which uses multi-head attention for pairs of proteins defined by the initial adjacency matrix. This only allows each protein to attend to its neighbors in the same genome subgraph. The output from the PST encoder are genome-contextualized protein embeddings, which are also the inputs to the PST decoder. The PST decoder uses multi-head attention pooling to project each contextualized protein embedding onto a learnable seed vector. This learns weights for each protein, which are used to pool each protein representation into a final genome representation. The full encoder-decoder PST shown here is framework used by the PST-TL (triplet loss) models, which output both contextualized protein embeddings and genome embeddings that are learned weighted averages of the preceding protein embeddings. Encoder-only PSTs that only have protein-level objectives like PST-MLM (masked language modeling) models stop after the second step, only natively outputting contextualized protein embeddings. Genome embeddings for encoder-only PSTs can be generated with a simple average over proteins for each genome.

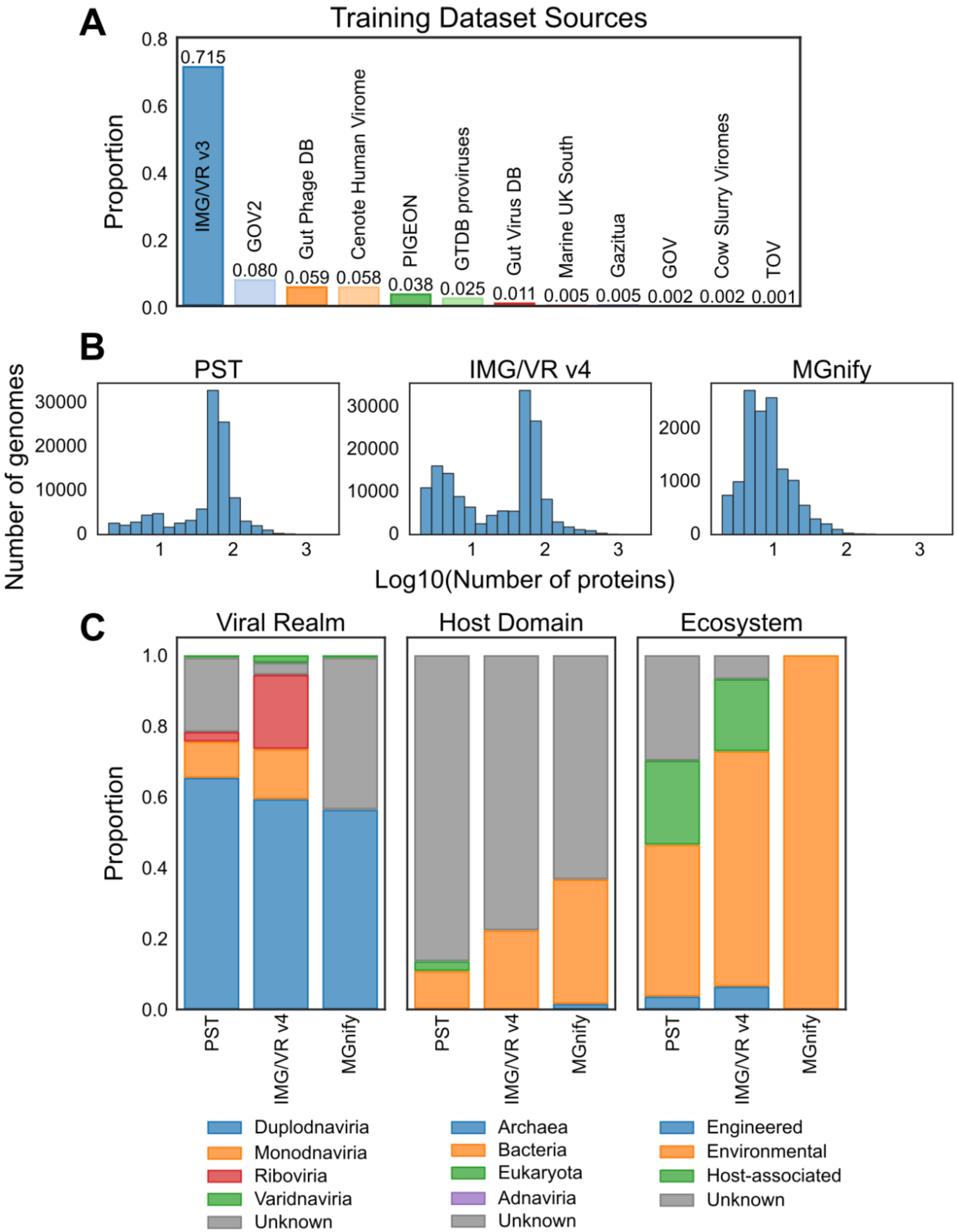


**Supplementary Figure 2**. Information about each dataset used for training and evaluation of PST. **A**) Proportional source of 103,589 training set viral genomes (see Supplemental Table 1 for the source publication of each virus). **B**) Distributions of the log10 number of proteins encoded per genome. **C**) The relative distributions of viral realm, host domain, and broad ecosystem for each dataset. Viral realm, if not provided by the source database, was predicted by geNomad. For the PST training and IMG/VR v4 datasets, host domains that were not provided by the source database, excluding predicted proviruses, were considered unknown. For the MGnify test dataset, putative hosts were predicted by iPHoP.

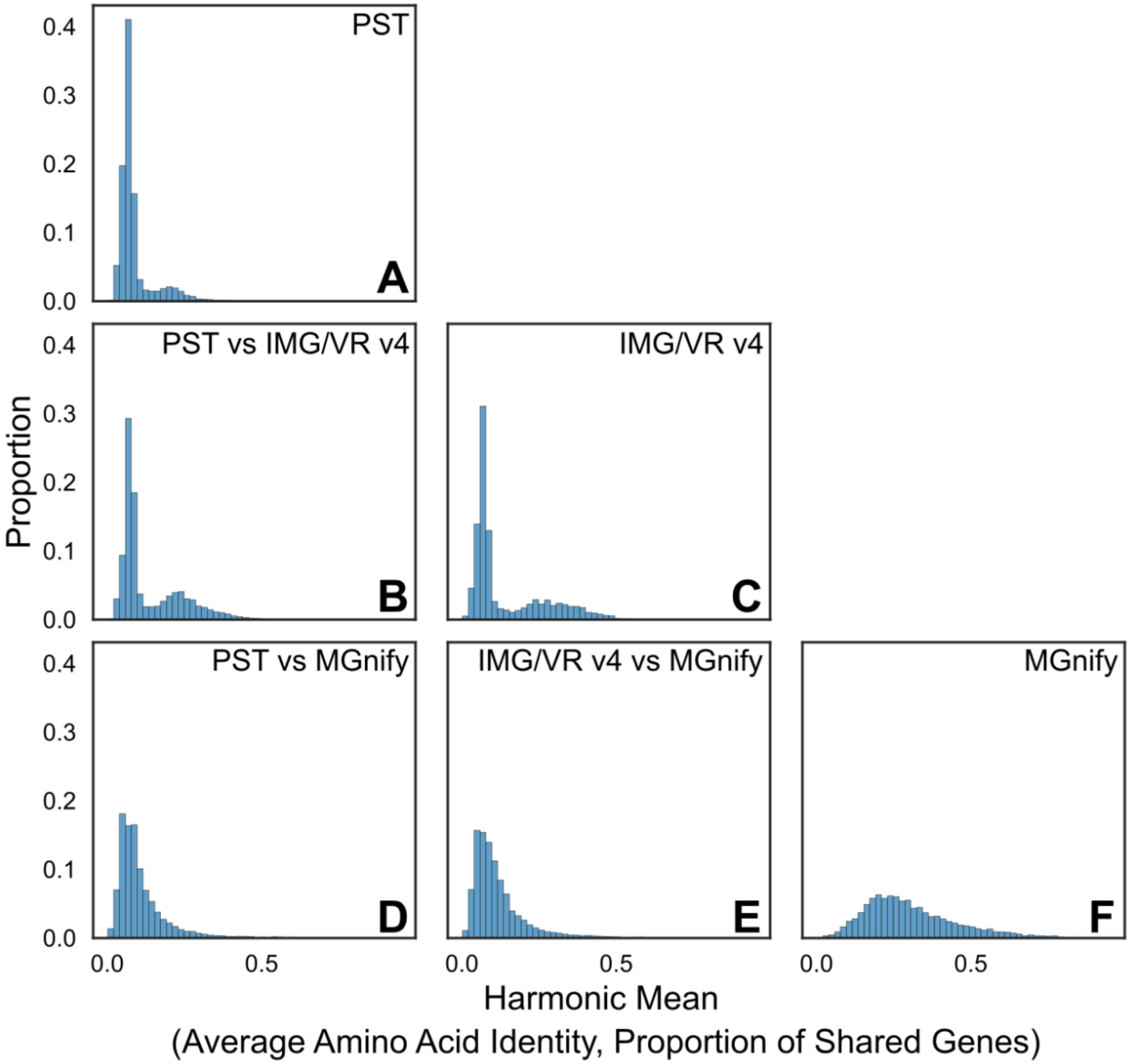


**Supplementary Figure 3**. Genome-genome similarity histograms comparing **A**) the PST training dataset with itself, **B**) the PST training dataset with the IMG/VR v4 test dataset, **C**) the IMG/VR v4 test dataset with itself, **D**) the PST training dataset with the MGnify test dataset, **E**) the IMG/VR v4 test dataset with the MGnify test dataset, and **F**) the MGnify test dataset with itself. Genome-genome similarity was computed as the harmonic mean of the Average Amino Acid Identity and the proportion of shared genes between each pair of genomes.

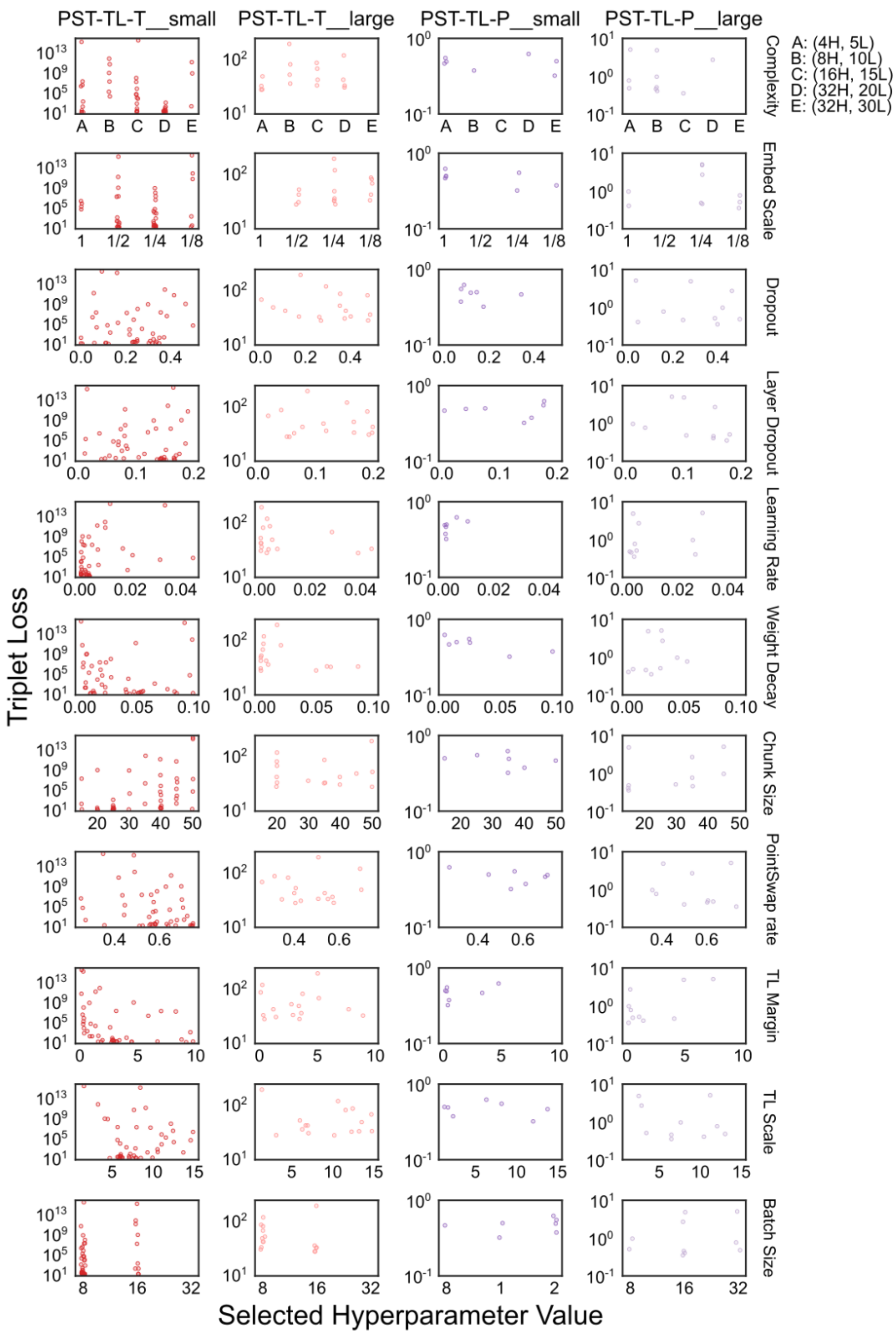


**Supplementary Figure 4**. Hyperparameter tuning for PST triplet loss models. Hyperparameters were tuned by cross validation based on taxonomic groups (T) or protein diversity groups (P). Rows indicate the specific hyperparameter, and columns indicate the model. There were 45 complete trials for PST-TL-T__small, 16 complete trials for PST-TL-T__large, 7 complete trials for PST-TL-P__small, and 10 complete trials for PST-TL-P__large. "Complexity" refers to the number of attention heads and encoder layers, with a key in the top right (H = attention heads, L = encoder layers, i.e. 4H = 4 attention heads, 5L = 5 encoder layers). "Embed scale" is the size of each of positional and strand embeddings relative to the input ESM2 protein embedding that are concatenated together. Weight decay is for the AdamW optimizer. The PST "chunk size" is the number of proteins per genome chunk. "PointSwap rate" is the proportion of proteins swapped between the anchor and positive genome during PointSwap sampling. “TL margin” is the minimum margin between anchor-positive and anchor-negative embedding distances before the indicated triplet no longer contributes to the loss. “TL scale” is the negative exponential decay scale factor to adjust the weight of the choice of the negative samples in the triplet loss function. Batch size is in units of number of genomes.

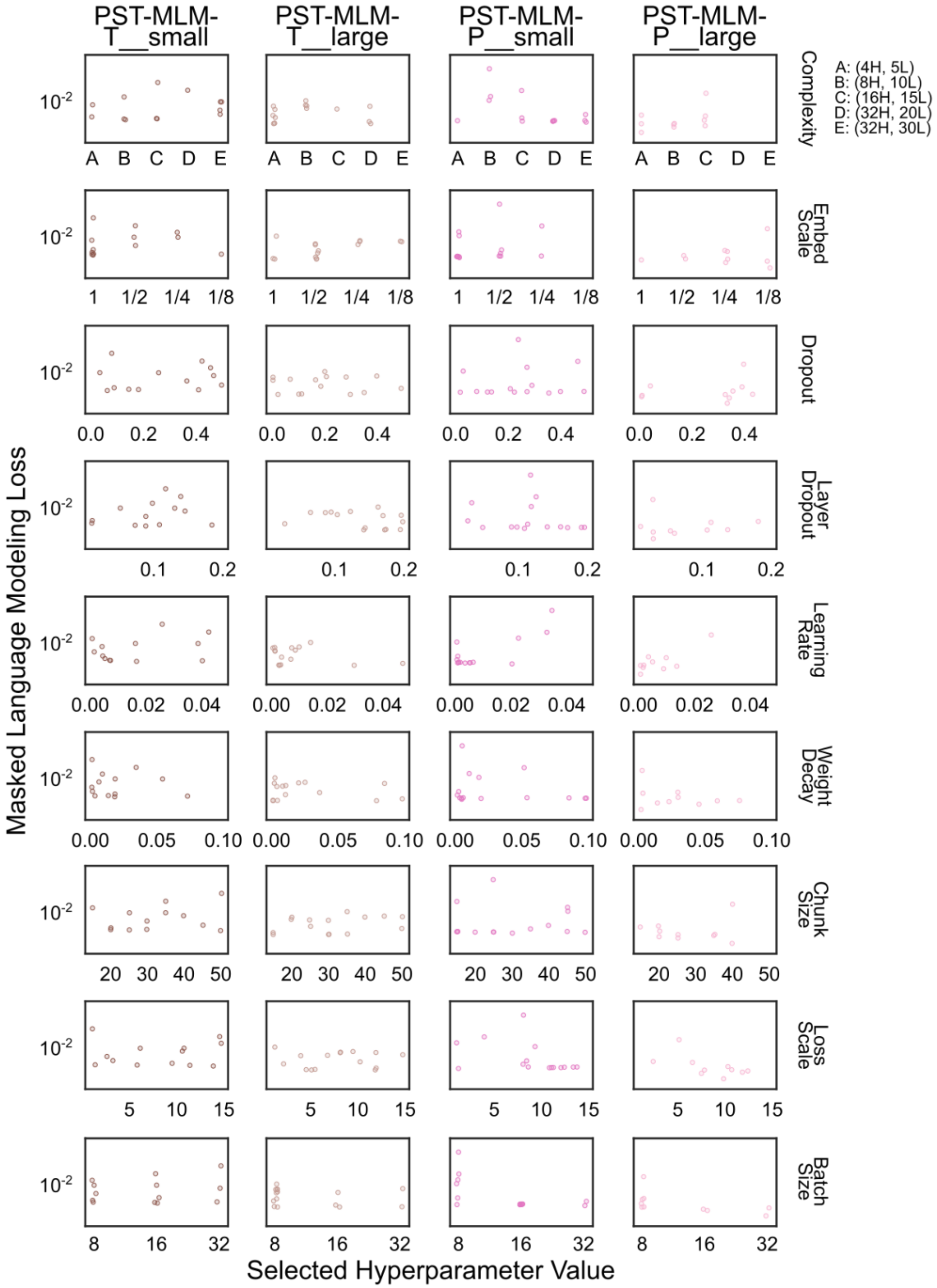


**Supplementary Figure 5**. Hyperparameter tuning for PST masked language modeling loss models. Hyperparameters were tuned by cross validation based on taxonomic groups (T) or protein diversity groups (P). Rows indicate the specific hyperparameter, and columns indicate the model. There were 13 complete trials for PST-MLM-T__small, 15 complete trials for PST-MLM-T__large, 15 complete trials for PST-MLM-P__small, and 10 complete trials for PST-MLM-P__large. "Complexity" refers to the number of attention heads and encoder layers, with a key in the top right (H = attention heads, L = encoder layers, i.e. 4H = 4 attention heads, 5L = 5 encoder layers). "Embed scale" is the size of each of positional and strand embeddings relative to the input ESM2 protein embedding that are concatenated together. Weight decay is for the AdamW optimizer. The PST "chunk size" is the number of proteins per genome chunk. "Loss scale" is the negative exponential decay scale factor to adjust the weight of the choice of the positive samples. Batch size is in units of number of genomes.

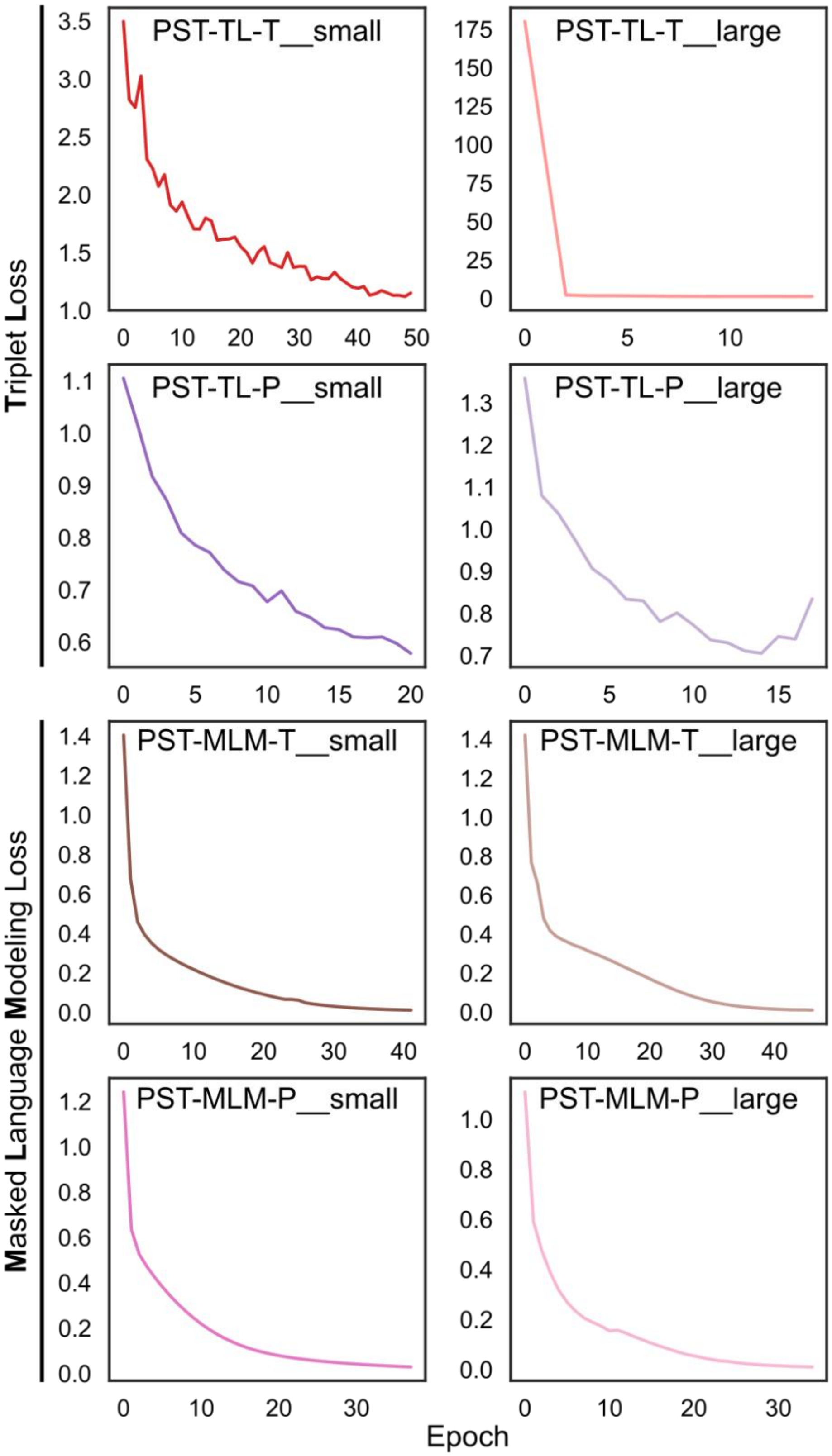


**Supplementary Figure 6**. Training loss curves for the final PST models after hyperparameters were chosen by cross validation. The names of each model indicate the loss objective (TL = Triplet Loss, MLM = Masked Language Modeling), how the cross validation groups were determined (T = viral taxonomic realm, P = protein diversity), and the size of the input ESM2 embeddings (small = esm2_t6_8M, large = esm2_t30_150M).

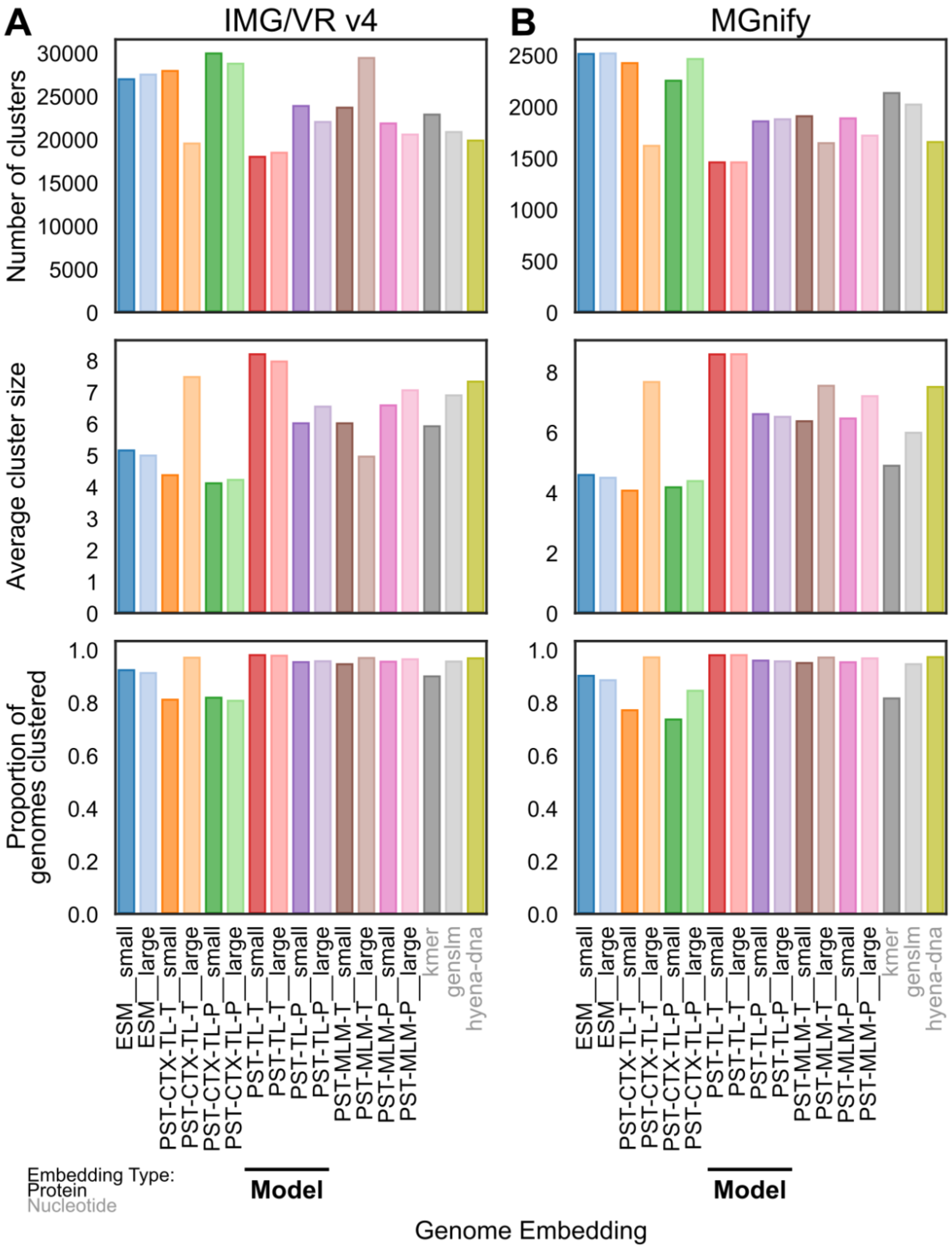


**Supplementary Figure 7**. Genome clustering stats for **A**) the IMG/VR v4 and **B**) MGnify test datasets. Genomes were clustered based on the angular similarity of L2-normalized genome embeddings from the corresponding embedding type on the x-axis. Singleton genomes not clustered were excluded from these stats.

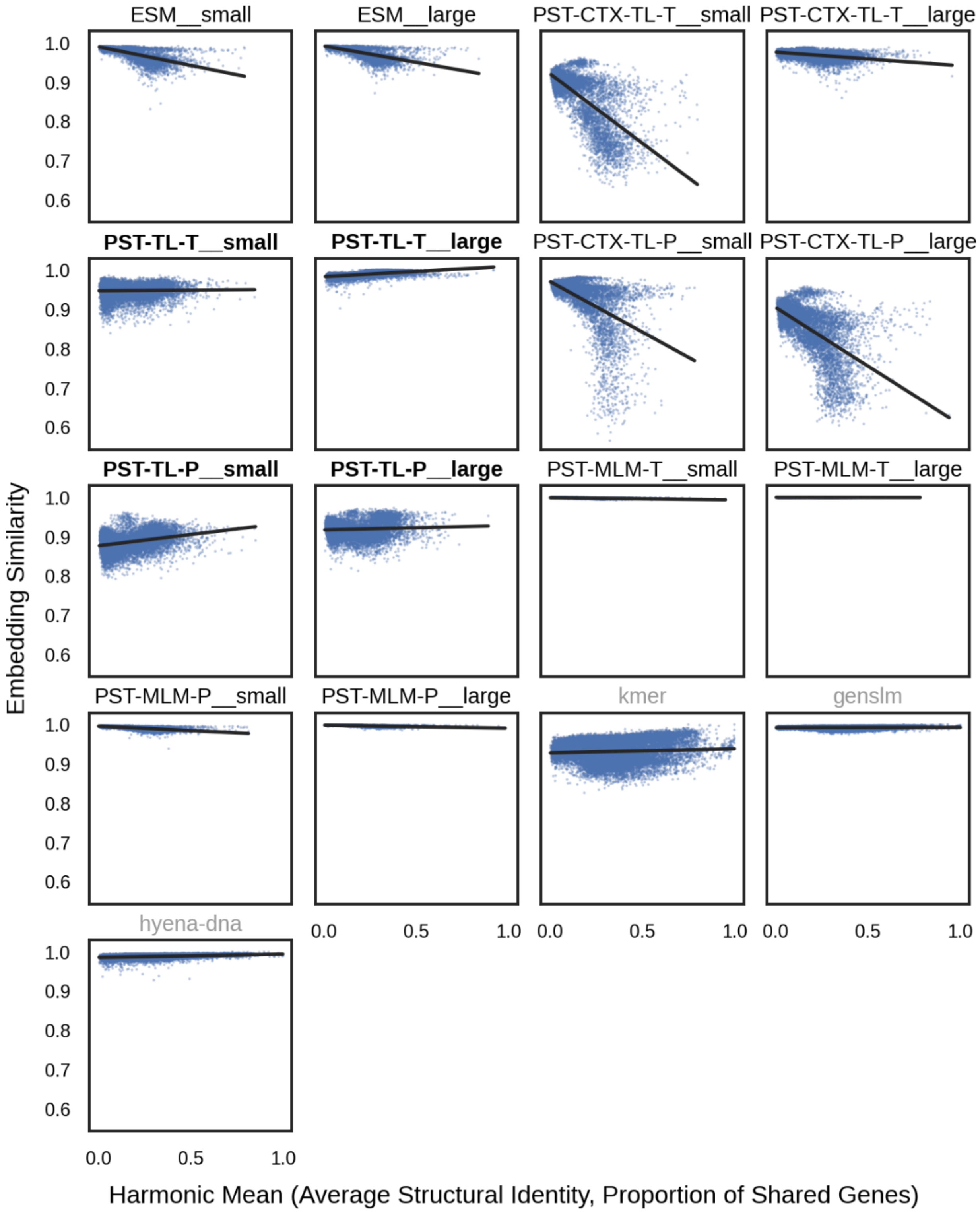


**Supplementary Figure 8**. Scatterplots of genome-genome similarity (x-axis) against embedding similarity (y-axis) for **closely related genomes** in the IMG/VR v4 test dataset. Genome-genome similarity is the harmonic mean of Average Structural Identity (ASI; see **Methods**) and the proportion of shared genes between each pair of genomes based on structural information. Each panel uses the corresponding genome embedding to compute the angular similarity between L2-normalized embeddings. ASI was only computed between genomes that clustered together based on the corresponding embedding. Genomes were defined as “similar” if there were any protein *sequence* alignments between the proteins from each pair of genomes.

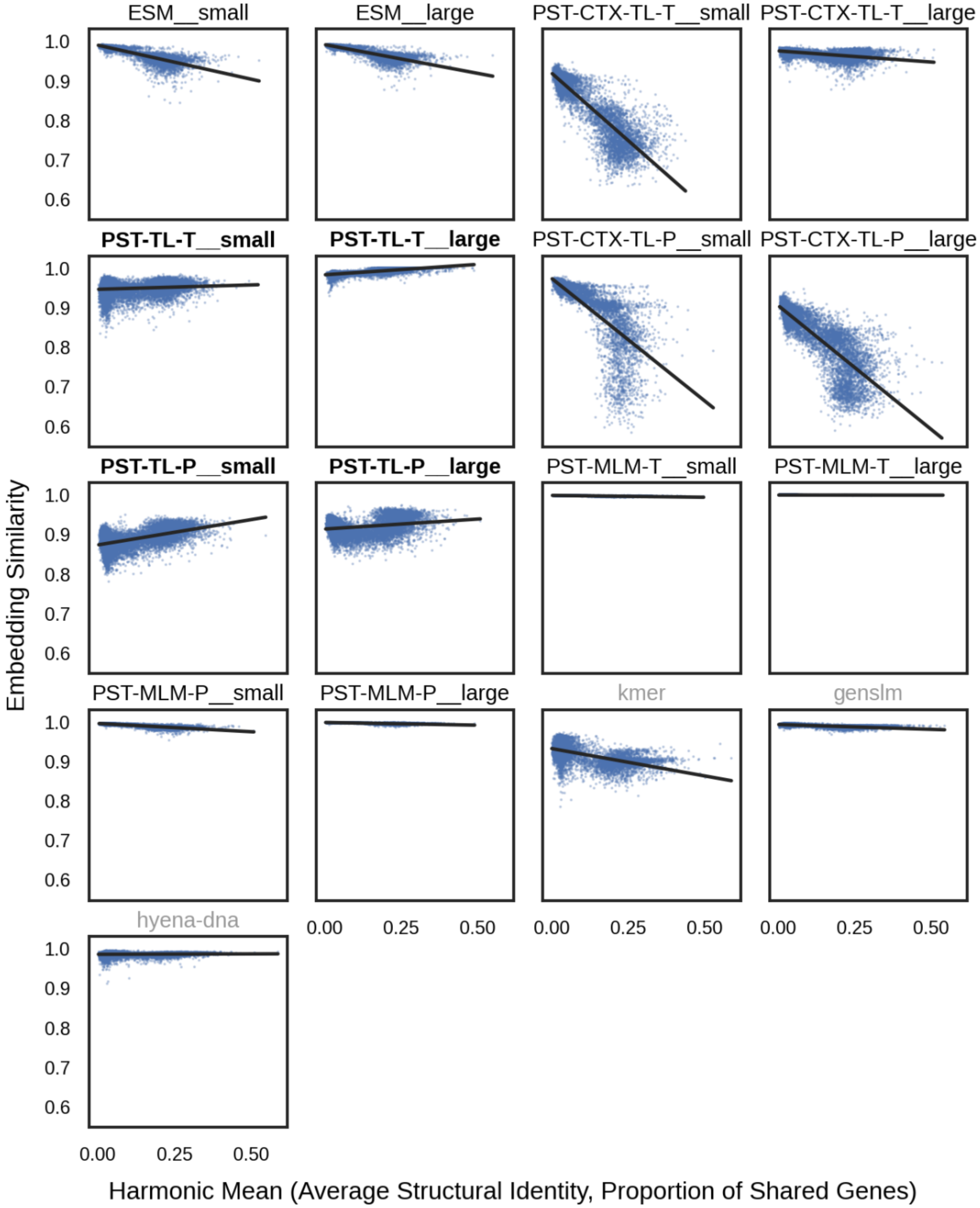


**Supplementary Figure 9**. Scatterplots of genome-genome similarity (x-axis) against embedding similarity (y-axis) for **distantly related genomes** in the IMG/VR v4 test dataset. Genome-genome similarity is the harmonic mean of Average Structural Identity (ASI; see **Methods**) and the proportion of shared genes between each pair of genomes based on structural information. Each panel uses the corresponding genome embedding to compute the angular similarity between L2-normalized embeddings. ASI was only computed between genomes that clustered together based on the corresponding embedding. Genomes were defined as “distant” if there were no protein *sequence* alignments and only protein *structural* alignments between the proteins from each pair of genomes.

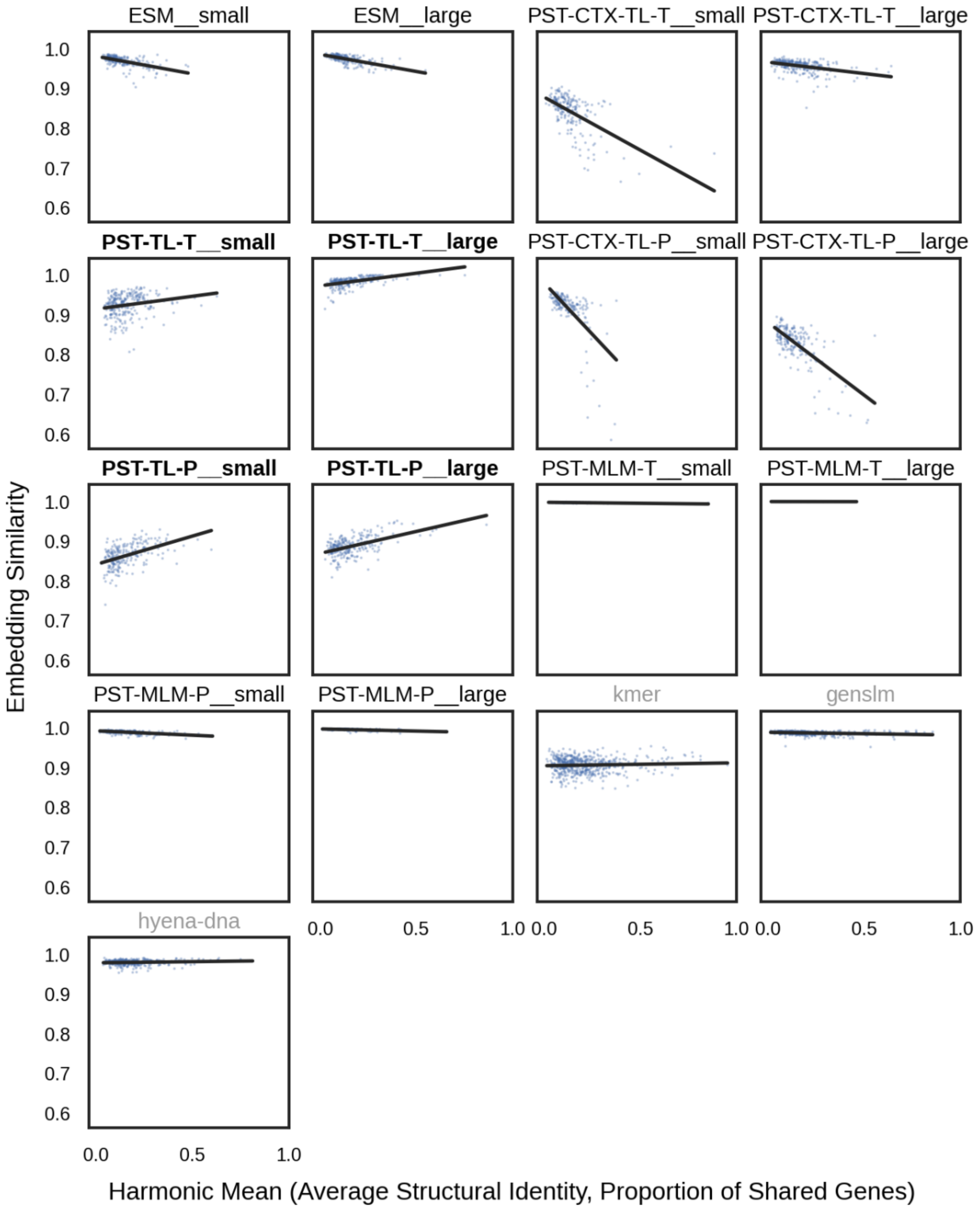


**Supplementary Figure 10**. Scatterplots of genome-genome similarity (x-axis) against embedding similarity (y-axis) for **closely related genomes** in the MGnify test dataset. Genome-genome similarity is the harmonic mean of Average Structural Identity (ASI; see **Methods**) and the proportion of shared genes between each pair of genomes based on structural information. Each panel uses the corresponding genome embedding to compute the angular similarity between L2-normalized embeddings. ASI was only computed between genomes that clustered together based on the corresponding embedding. Genomes were defined as “similar” if there were any protein *sequence* alignments between the proteins from each pair of genomes.

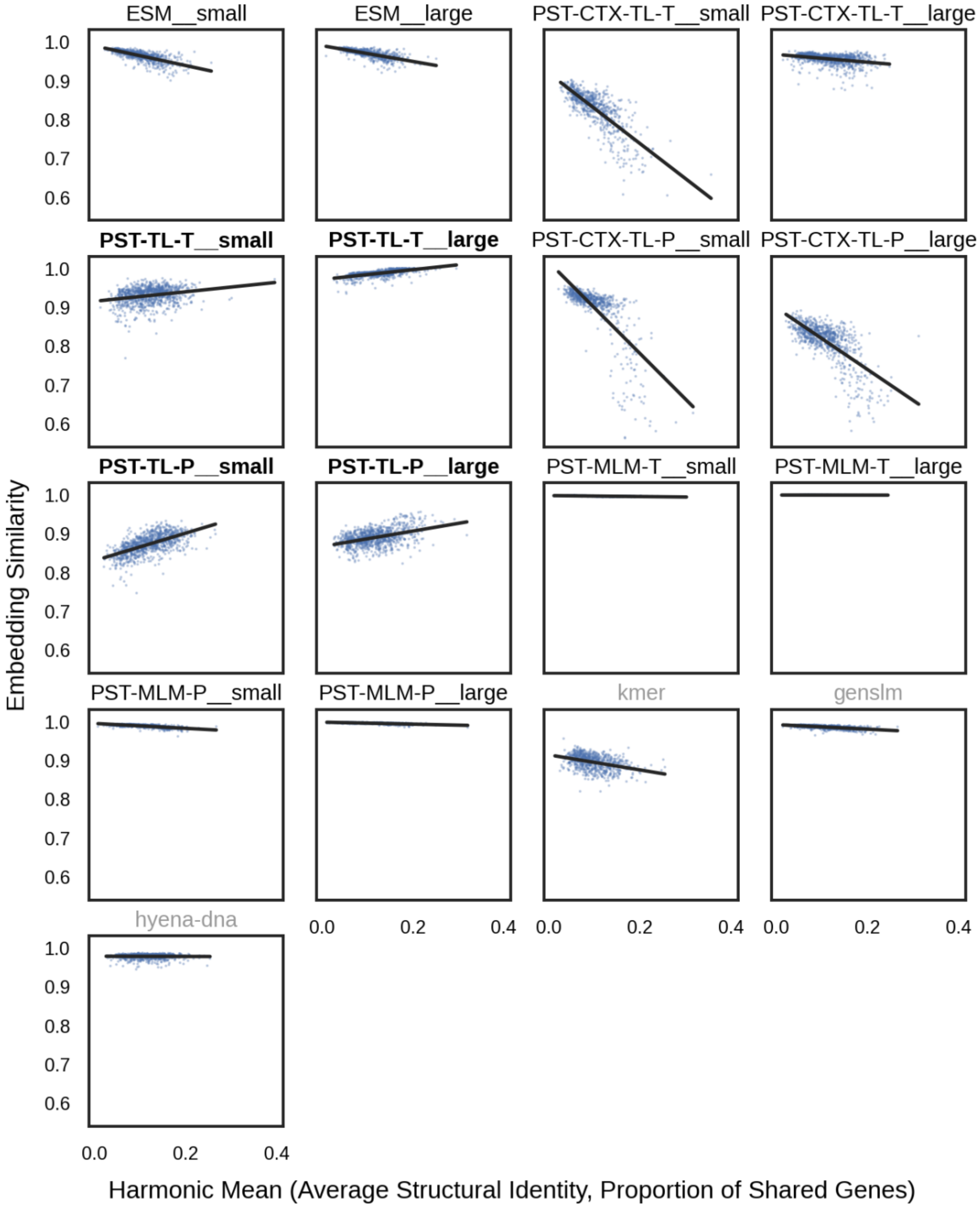


**Supplementary Figure 11**. Scatterplots of genome-genome similarity (x-axis) against embedding similarity (y-axis) for **distantly related genomes** in the MGnify test dataset. Genome-genome similarity is the harmonic mean of Average Structural Identity (ASI; see **Methods**) and the proportion of shared genes between each pair of genomes based on structural information. Each panel uses the corresponding genome embedding to compute the angular similarity between L2-normalized embeddings. ASI was only computed between genomes that clustered together based on the corresponding embedding. Genomes were defined as “distant” if there were no protein *sequence* alignments and only protein *structural* alignments between the proteins from each pair of genomes.

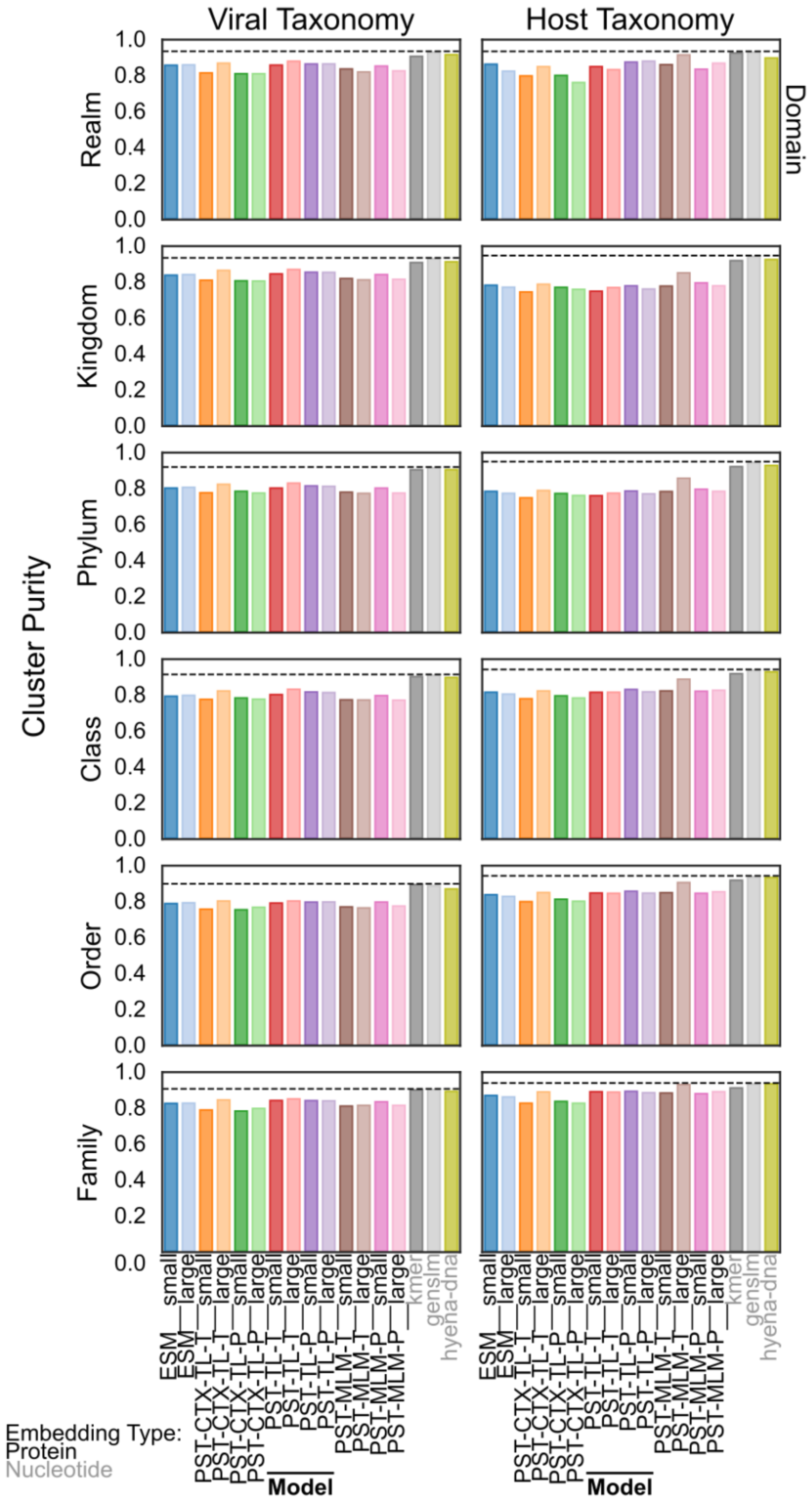


**Supplementary Figure 12**. Viral and Host taxonomic purity for the IMG/VR v4 test dataset faceted by taxonomic rank. Purity is defined as the cluster size-weighted average of information gain ratio for non-singleton genome clusters. Missing or unknown taxonomic labels were excluded in clusters that had at least 1 labeled genome.

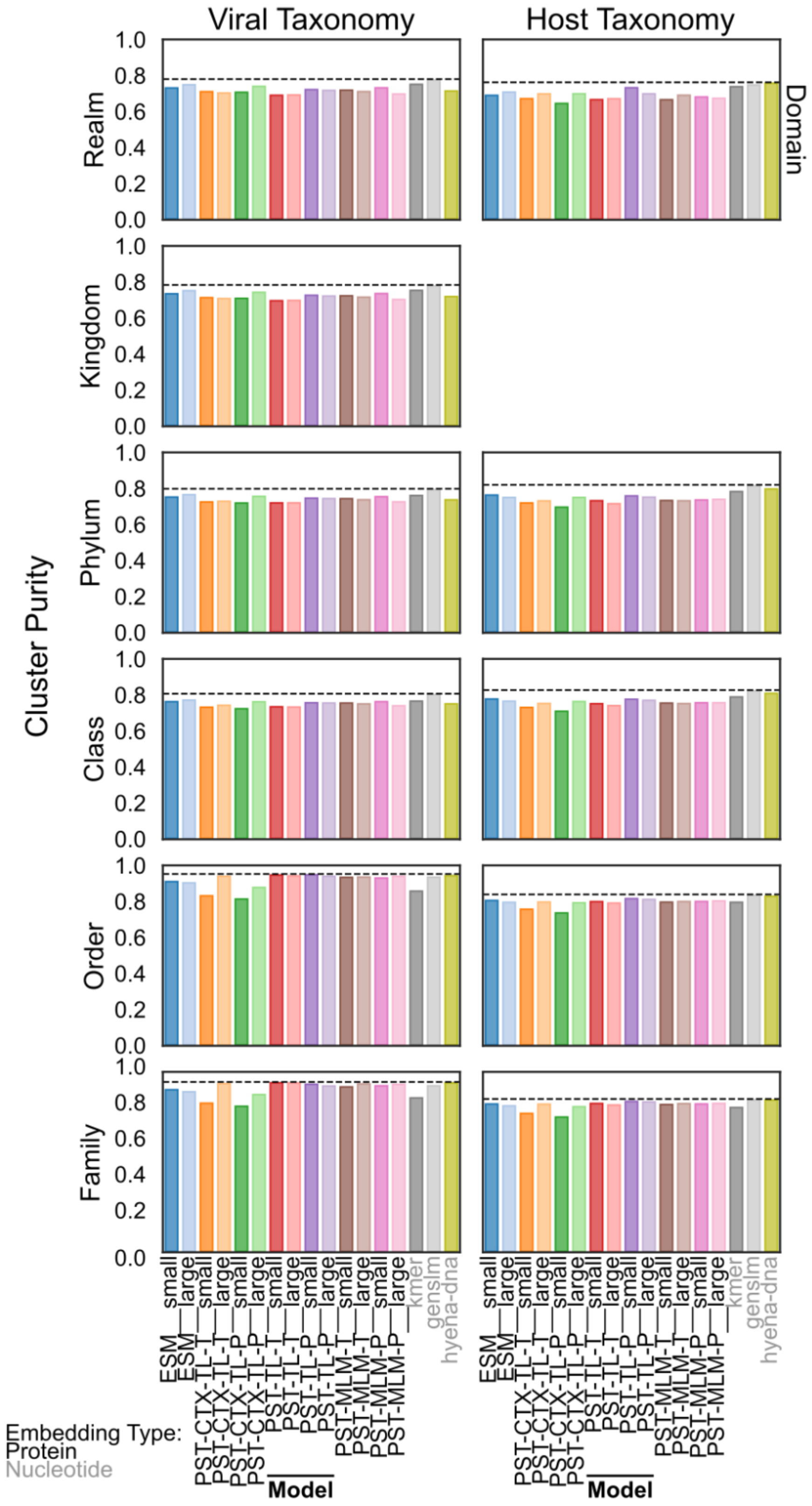


**Supplementary Figure 13**. Viral and Host taxonomic purity for the MGnify test dataset faceted by taxonomic rank. Purity is defined as the cluster size-weighted average of information gain ratio for non-singleton genome clusters. Missing or unknown taxonomic labels were excluded in clusters that had at least 1 labeled genome. Host taxonomic labels were predicted by iPHoP, which does not output a predicted host kingdom even when it could be inferred from the domain.

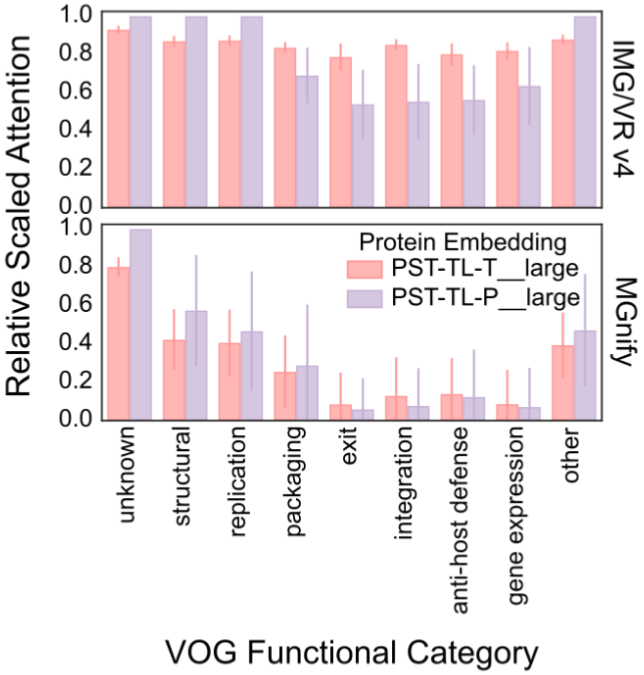


**Supplementary Figure 14**. Relative scaled attention from large triplet loss PSTs for proteins belonging to different VOG functional categories for IMG/VR v4 (top) and MGnify (bottom) test datasets. The per-protein attention scores from each model were normalized by the number of proteins encoded per scaffold (see **Methods**). For each model, these normalized attention values were then rescaled so that the maximum value was 1. Then the top 1,000 proteins based on the rescaled normalized attention were chosen for each functional category. The bar height represents the mean of those attention values, and the bars indicate ±1 standard deviation.

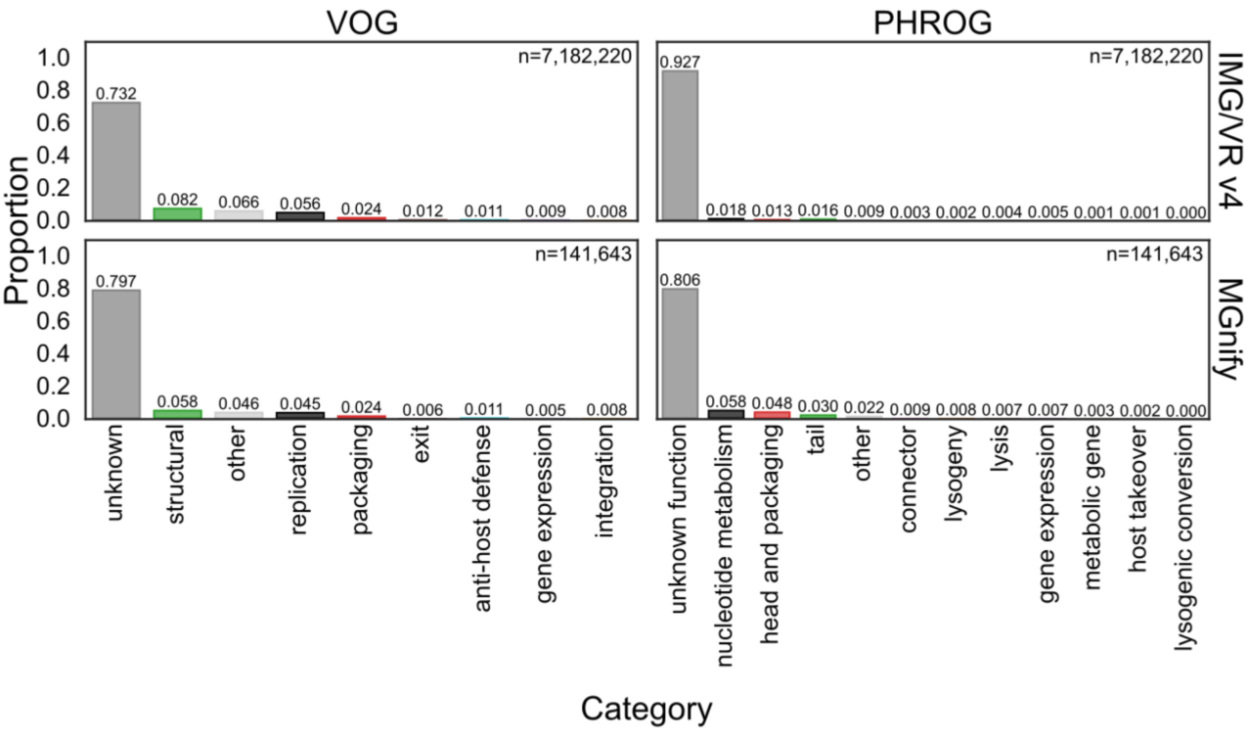


**Supplementary Figure 15**. Distribution of protein functional annotations using curated categories from the VOG (left) or PHROG (right) databases for the proteins from the IMG/VR v4 (top) and MGnify (bottom) test datasets.

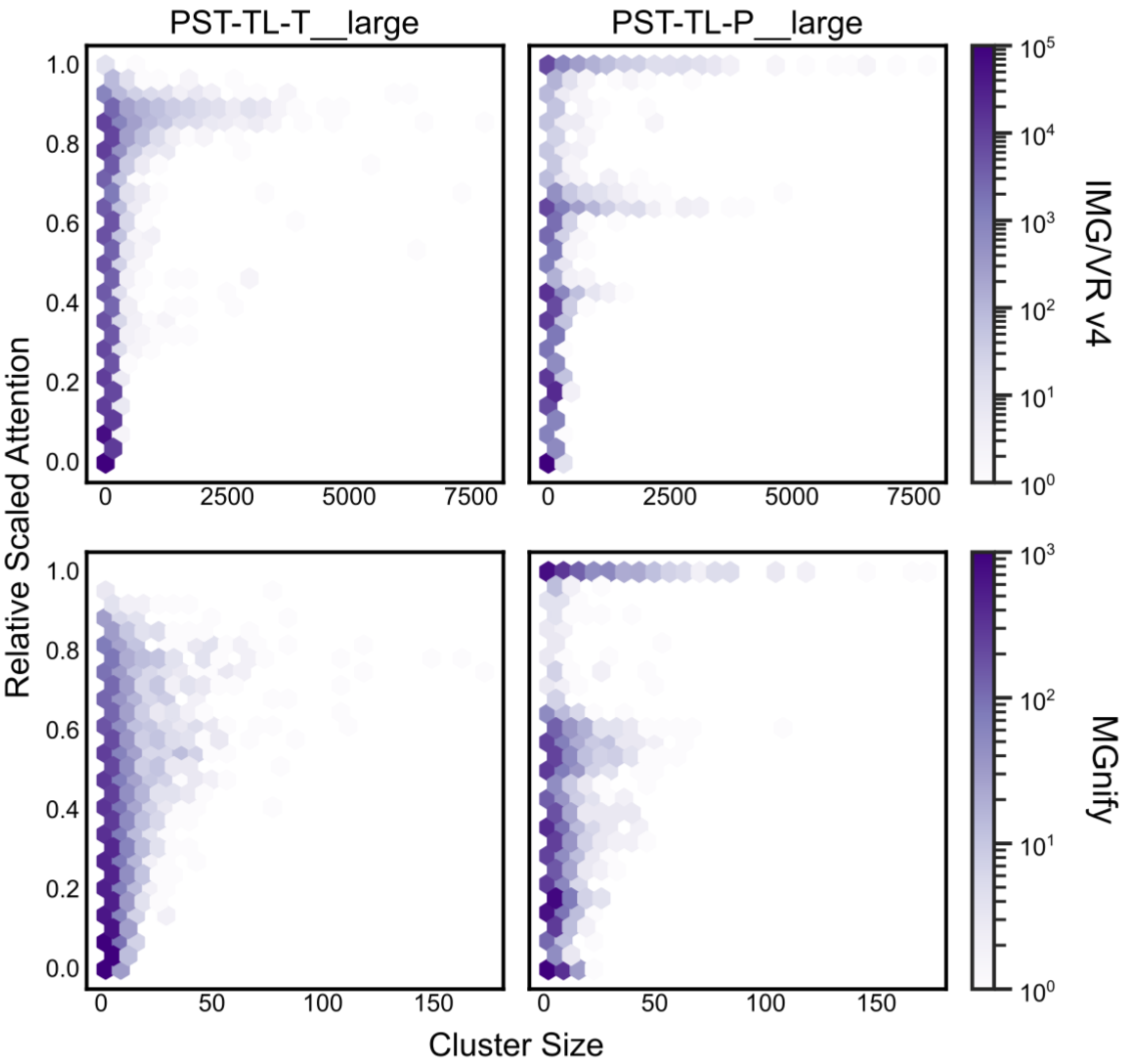


**Supplementary Figure 16**. Relative scaled attention from large triplet loss PSTs compared against the size of sequence identity-based clusters. The per-protein attention scores from each model were normalized by the number of proteins encoded per scaffold (see **Methods**). For each model, these normalized attention values were then rescaled so that the maximum value was 1. Then, for each protein cluster, the maximum scaled normalized attention was selected for the hexagonal histograms. The shade indicates the number of proteins in each bin (log scale).

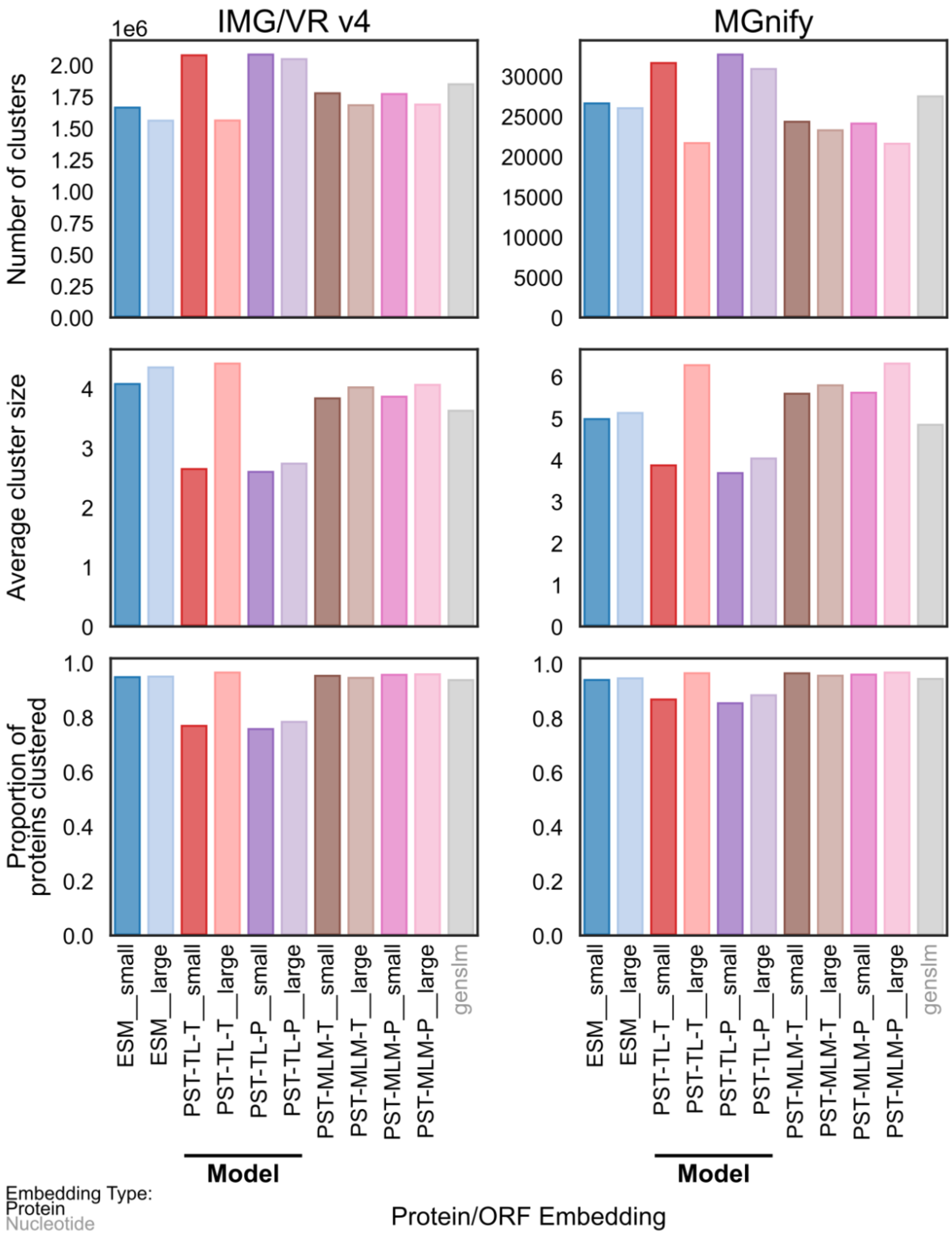


**Supplementary Figure 17**. Protein clustering stats for the IMG/VR v4 and MGnify test datasets. Proteins were clustered based on the angular similarity of L2-normalized protein embeddings from the corresponding embedding type on the x-axis. Singleton proteins not clustered were excluded from these stats.

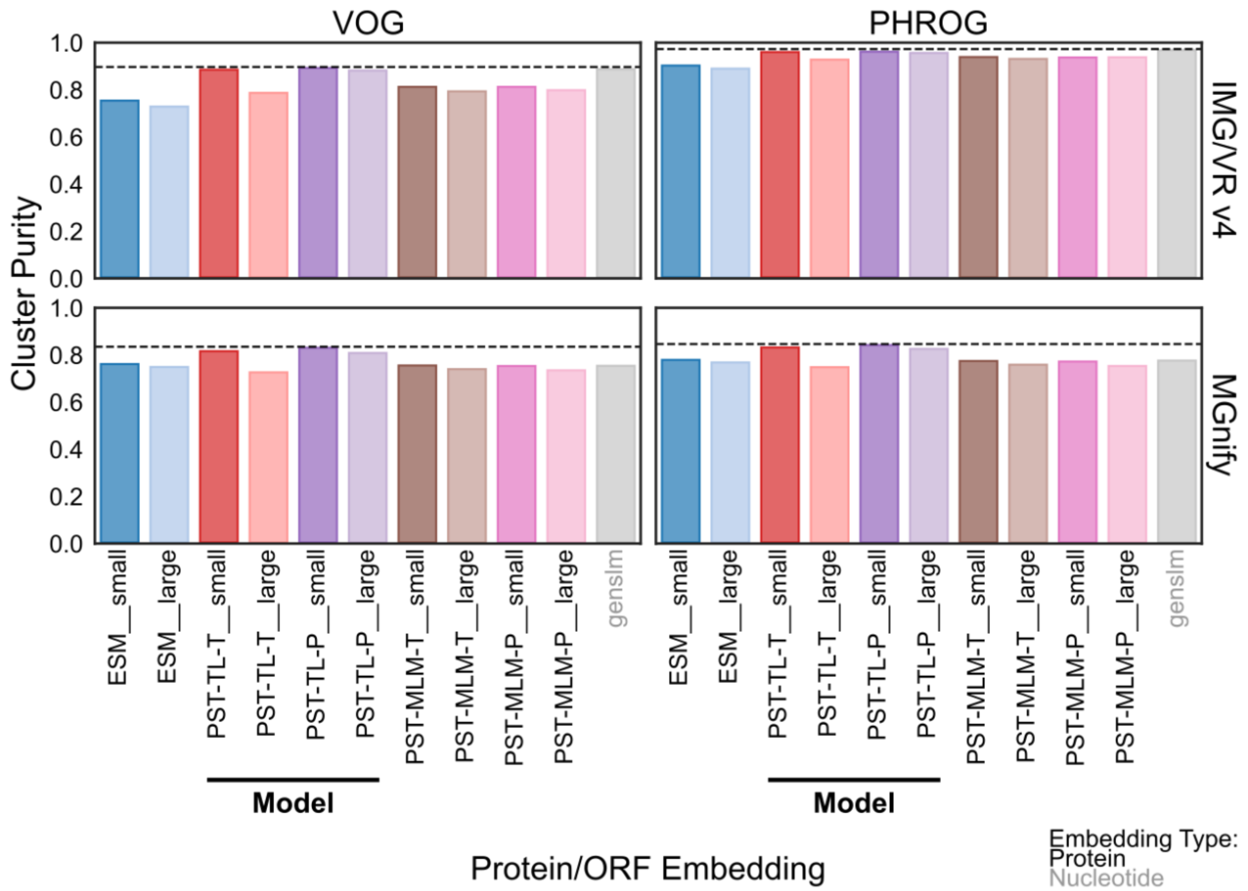


**Supplementary Figure 18**. Protein cluster functional purity for the IMG/VR v4 (top) and MGnify (bottom) test datasets when annotated with the VOG (left) or PHROG (right) databases. Proteins were clustered based on the angular similarity of L2-normalized protein embeddings from the corresponding embedding type on the x-axis. Only clusters that had at least 1 labeled protein were included, and in these clusters, all proteins without a predicted function were ignored. Purity is defined as the cluster size-weighted average of information gain ratio for these protein clusters.

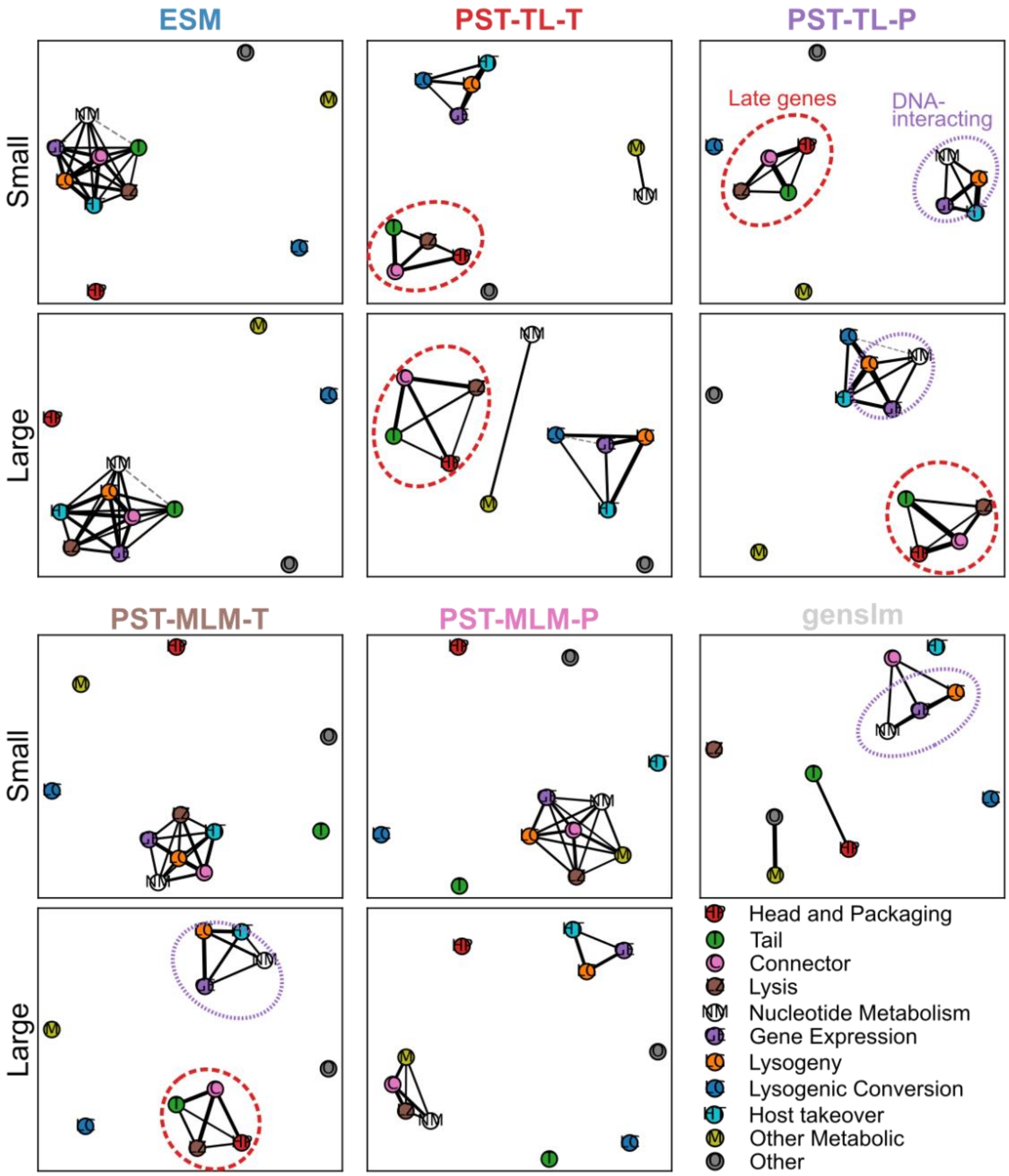


**Supplementary Figure 19**. Protein function co-clustering summary for the IMG/VR v4 test dataset based on PHROG annotations. Rows indicate the model size (i.e. the first panel is "ESM__small", excluding GenSLM), and columns indicate the protein embedding type. Each connected component was determined by Leiden community detection in co-clustering graph with a resolution of 1.25 (see **Methods**). Edges indicate functional categories that were more enriched in protein clusters compared to the joint occurrence of these categories in the PHROG database. The length of the edges inversely indicate the degree of enrichment (i.e. shorter edge = more enriched). Dotted gray edges indicate connections that were less enriched than expected but still clustered together. Red dashed circles highlight detected late gene modules, composed of Head and Packaging, Tail, Connector, and Lysis. Purple dotted circles indicate detected DNA-interacting modules, consisting of Nucleotide Metabolism, Gene Expression, and Lysogeny.

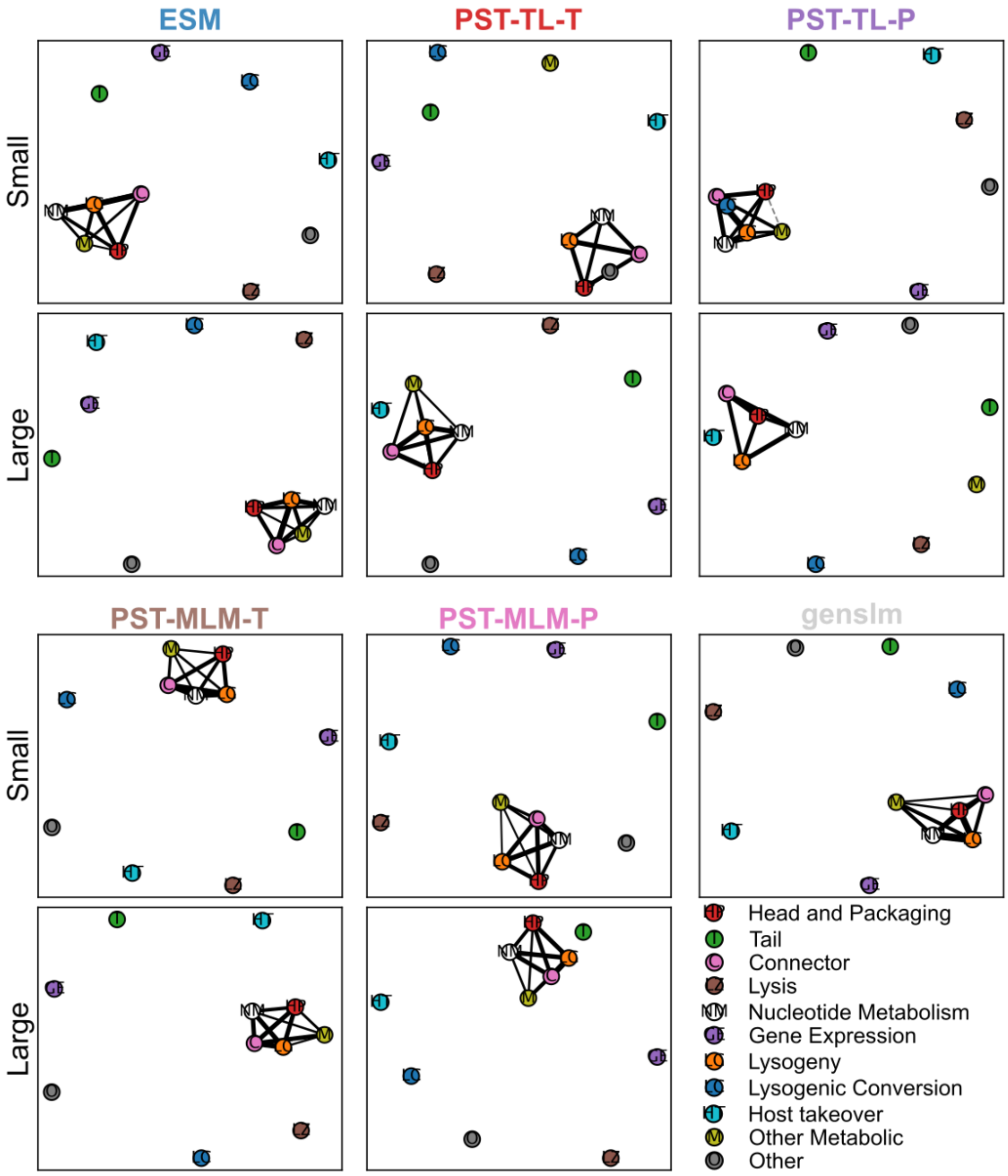


**Supplementary Figure 20**. Protein function co-clustering summary for the MGnify test dataset based on PHROG annotations. Rows indicate the model size (i.e. the first panel is "ESM__small", excluding GenSLM), and columns indicate the protein embedding type. Each connected component was determined by Leiden community detection in co-clustering graph with a resolution of 1.25 (see **Methods**). Edges indicate functional categories that were more enriched in protein clusters compared to the joint occurrence of these categories in the PHROG database. The length of the edges inversely indicate the degree of enrichment (i.e. shorter edge = more enriched). Dotted gray edges indicate connections that were less enriched than expected but still clustered together.

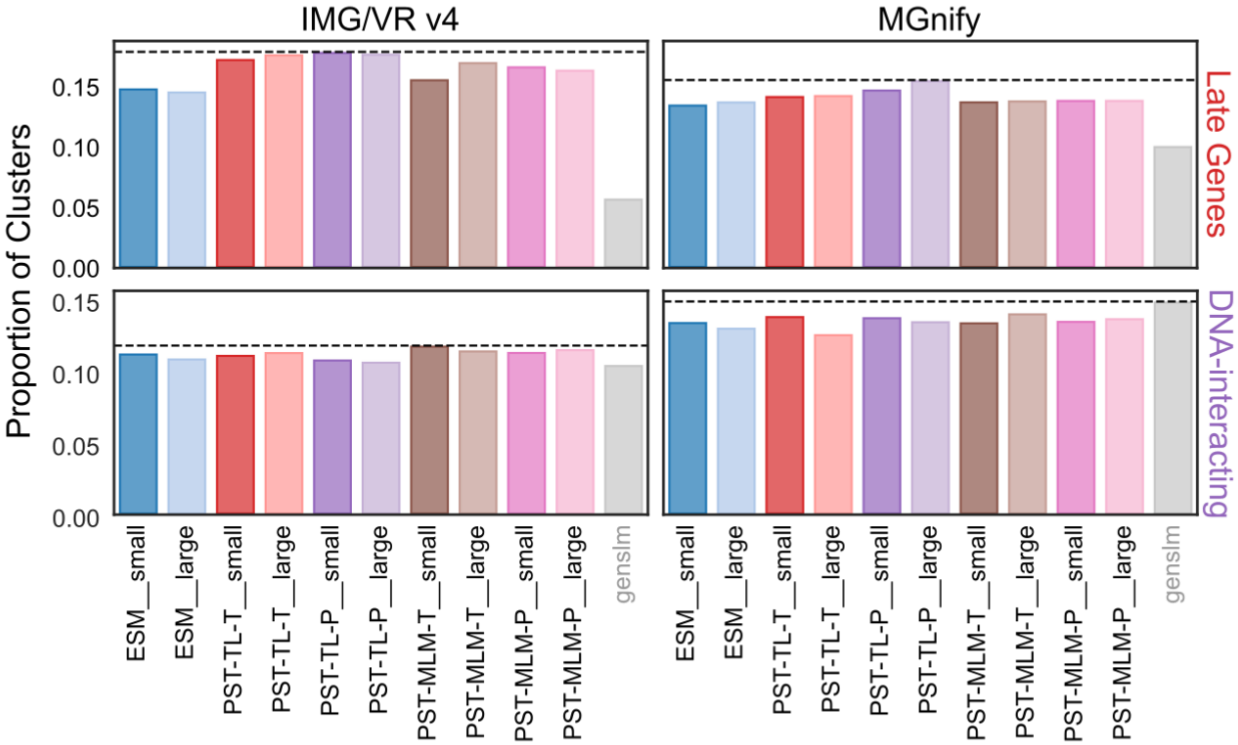


**Supplementary Figure 21**. The proportion of protein clusters that correspond to Late Gene (top) or DNA-interacting (bottom) modules for the IMG/VR v4 (left) and MGnify (right) test datasets. A protein cluster was considered to represent a functional module if all proteins annotated by VOG in the cluster belonged to the subcategories that composed each module (Late Genes: Structural, Exit, Packaging; DNA-interacting: Replication, Integration, Packaging, Gene Expression). Each protein cluster was also required to have annotated proteins belonging to at least 2 of the subcategories.

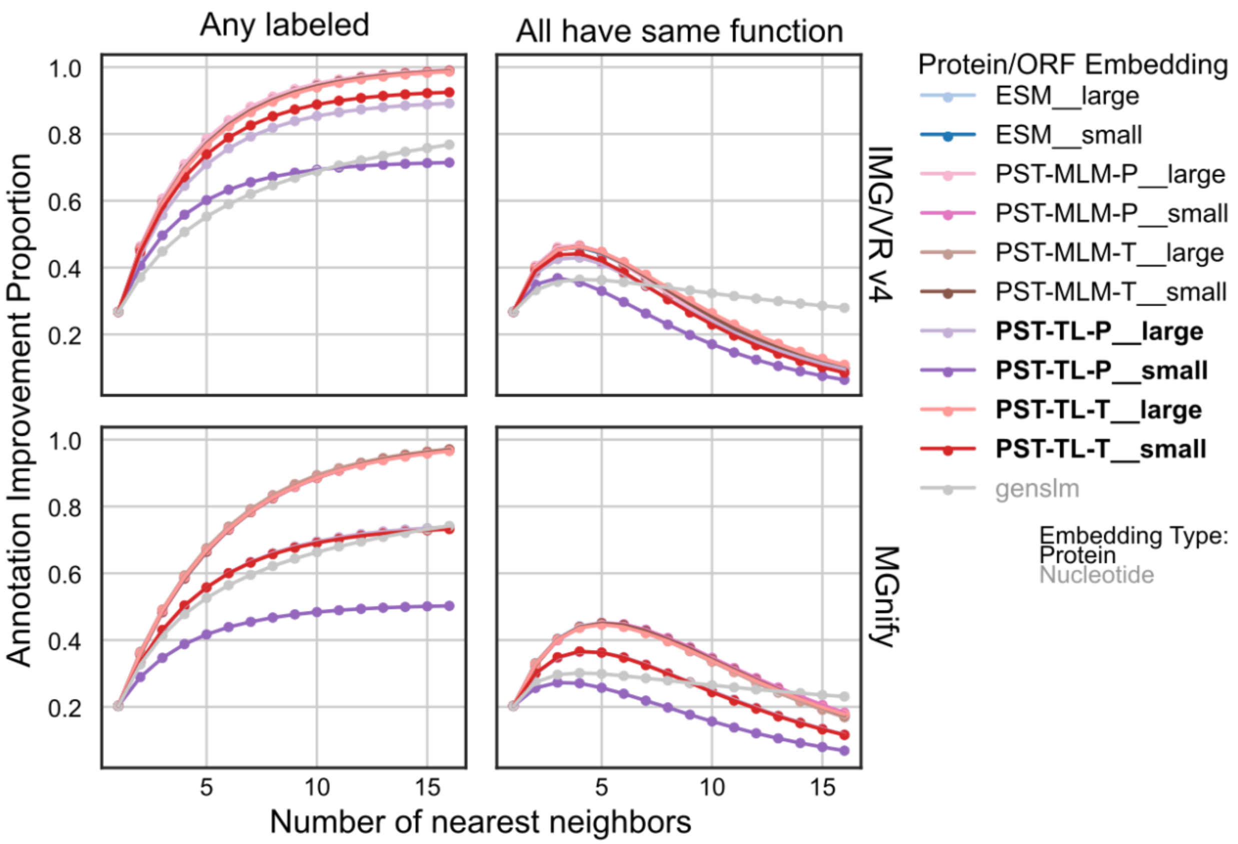


**Supplementary Figure 22**. The proportion of proteins unannotated by VOG whose nearest neighbors in embedding space are annotated. The colors indicate the protein/ORF embedding. Nearest neighbors were searched after L2-normalizing the protein embeddings using angular similarity. Left: A hit was considered for each unannotated protein if any of the neighbors less than or equal to the current number of nearest neighbors were annotated. Right: A hit was considered similarly to the left side with the additional constraint that all of the current set of nearest neighbors must belong to the same VOG functional category. Unannotated proteins were not used to penalize the score.

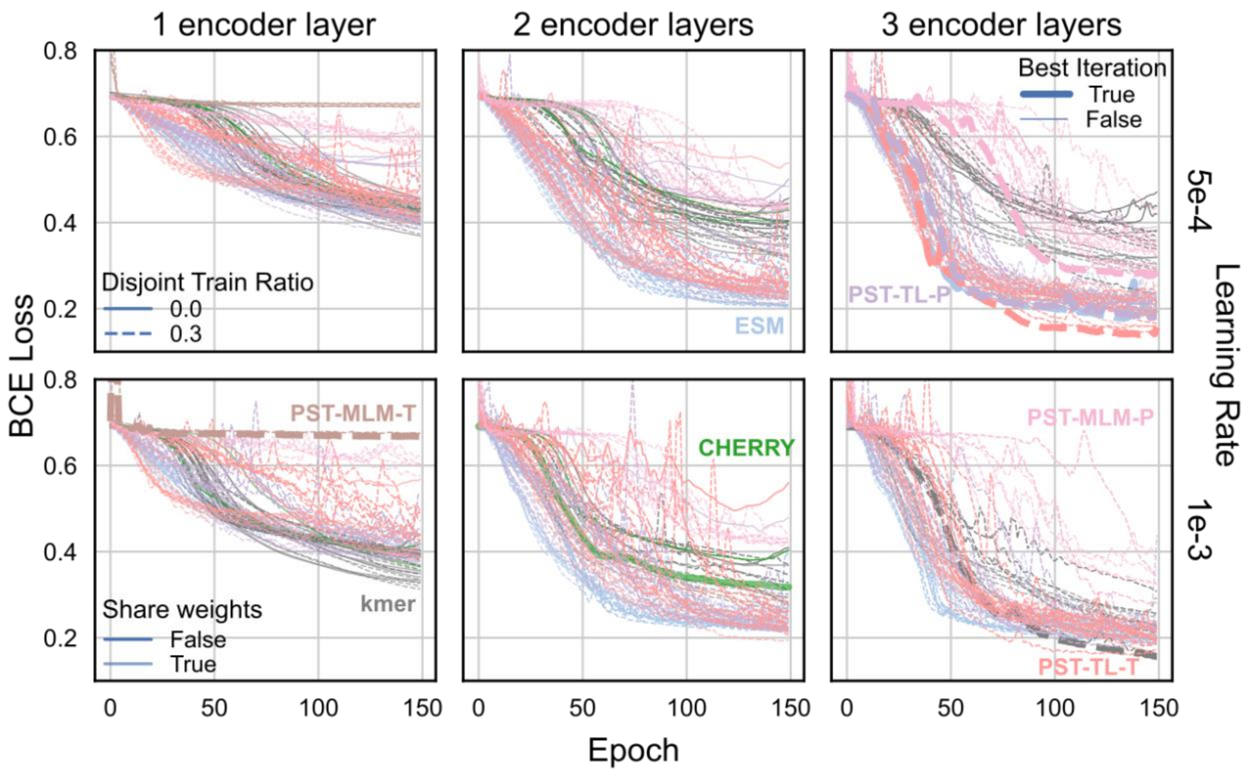


**Supplementary Figure 23**. Binary cross entropy loss curves for graph-based host prediction models. Columns indicate the number of encoder layers, and rows indicate the learning rate for the AdamW optimizer. Colors indicate the genome embedding used for the node embeddings of both virus and host genomes. For all embedding types except CHERRY and kmer, the indicated genome embedding was also used to cluster viruses to create virus-virus edges. Bold lines indicate the model iteration that was chosen based on the minimum loss during the final 20 epochs. Solid / dashed lines indicate the disjoint training ratio, which allocates a certain proportion of edges for a validation set during training. In the case of 0.0, all training edges were used for message passing and inference. The opacity of the lines indicate if the model weights were shared between the virus-host and virus-virus edges during message passing in the same layer. Only curves with a Spearman correlation ≤-0.9 were included to select iterations where the loss was strongly monotonically decreasing.

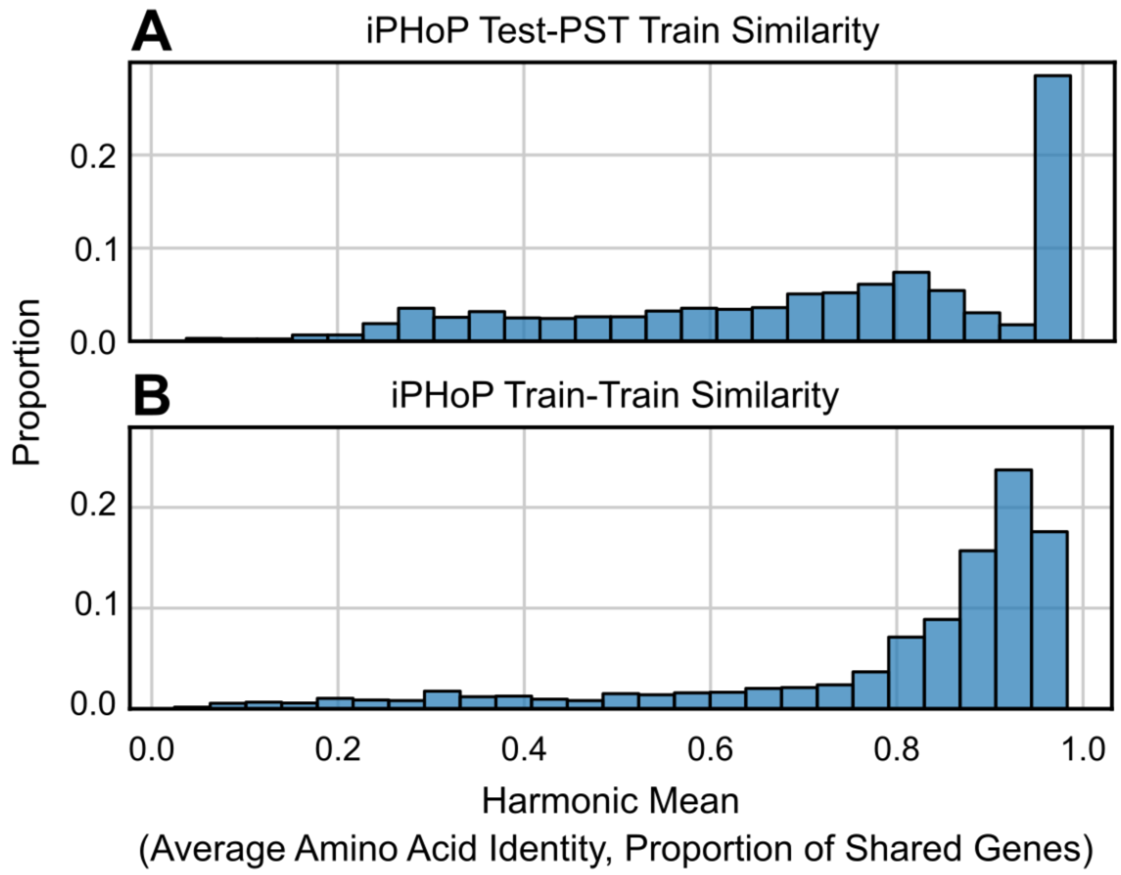


**Supplementary Figure 24**. Genome-genome similarity between viruses from the indicated datasets. Genome-genome similarity was computed as the harmonic mean of the Average Amino Acid Identity and the proportion of shared genes between each pair of genomes. Then, the maximum similarity score for each iPHoP test virus when searching against the **A**) PST training dataset or **B**) same dataset was kept.

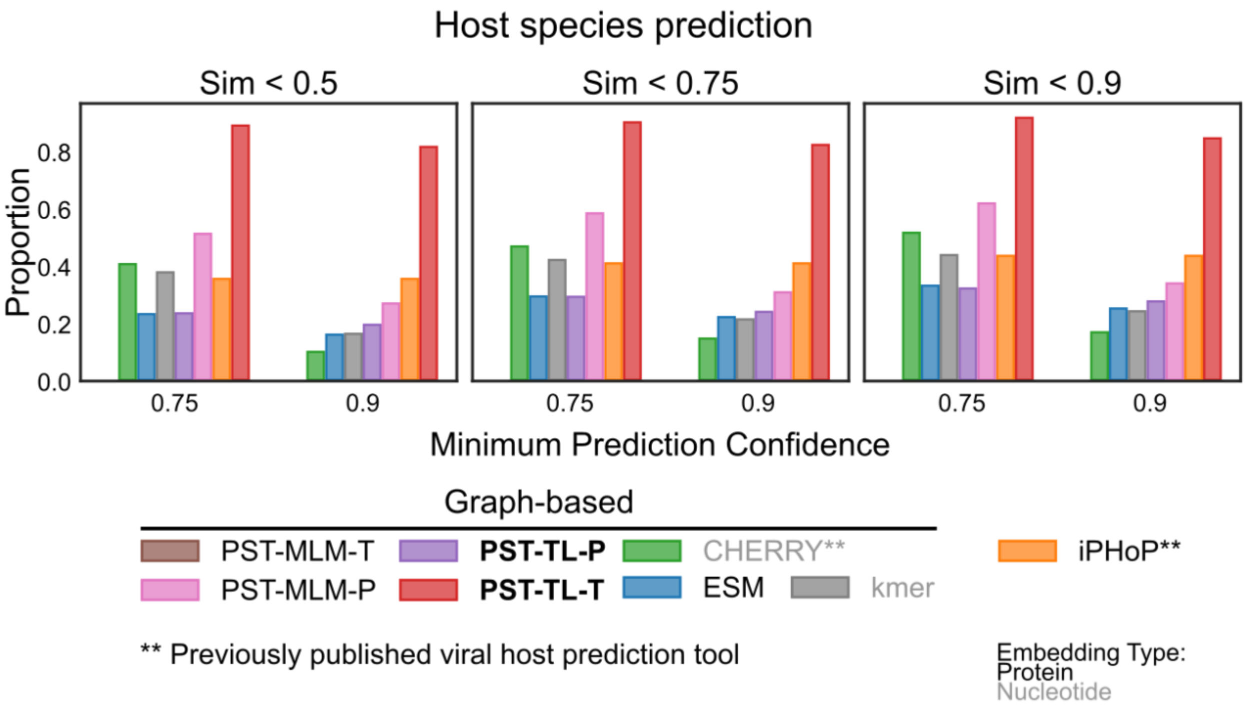


**Supplementary Figure 25**. Host species prediction performance removing viruses that were similar to the PST training dataset. Columns indicate the similarity threshold based on the harmonic mean of the Average Amino Acid Identity and the proportion of shared genes between each pair of genomes when comparing the iPHoP test and PST training datasets. Colors indicate the genome embedding type used to represent both the virus and host genomes in the interaction graph. The x-axis indicates the minimum prediction confidence required, and PST-MLM-T had no predictions above these thresholds.

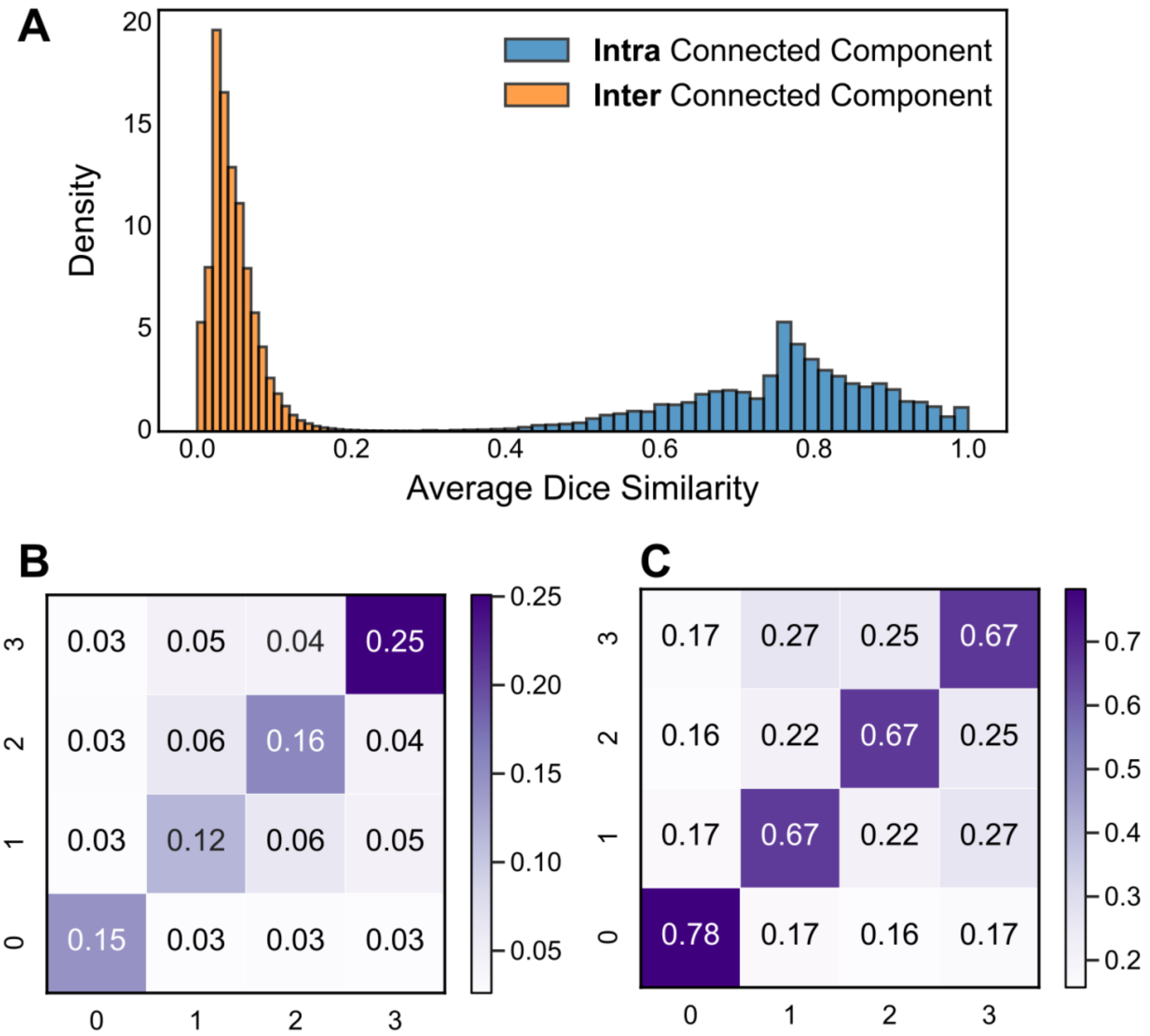


**Supplementary Figure 26**. Evaluation of 4 cross validation (CV) groups based on separating genomes into groups with minimally overlapping shared protein content. **A**) The average dice score of all pairs of genomes in the same connected component (blue) and across connected components (orange) was plotted as a histogram. Dice scores were computed using protein cluster presence/absence binary matrices for all genomes (see **Methods**). **B**) The average dice score between each of the 4 largest connected components used as seeds to expand each protein diversity CV group. **C**) The average maximum dice score between each genome and all others from each of the 4 protein diversity CV groups.
